# *Wolbachia*-induced inhibition of O’nyong nyong virus in *Anopheles* mosquitoes is mediated by Toll signaling and modulated by cholesterol

**DOI:** 10.1101/2023.05.31.543096

**Authors:** Sujit Pujhari, Grant L Hughes, Nazzy Pakpour, Yasutsugu Suzuki, Jason L Rasgon

## Abstract

Enhanced host immunity and competition for metabolic resources are two main competing hypotheses for the mechanism of *Wolbachia*-mediated pathogen inhibition in arthropods. Using an *Anopheles* mosquito – somatic *Wolbachia* infection – O’nyong nyong virus (ONNV) model, we demonstrate that the mechanism underpinning *Wolbachia*-mediated virus inhibition is up-regulation of the Toll innate immune pathway. However, the viral inhibitory properties of *Wolbachia* were abolished by cholesterol supplementation. This result was due to *Wolbachia*-dependent cholesterol-mediated suppression of Toll signaling rather than competition for cholesterol between *Wolbachia* and virus. The inhibitory effect of cholesterol was specific to *Wolbachia*-infected *Anopheles* mosquitoes and cells. These data indicate that both *Wolbachia* and cholesterol influence Toll immune signaling in *Anopheles* mosquitoes in a complex manner and provide a functional link between the host immunity and metabolic competition hypotheses for explaining *Wolbachia*-mediated pathogen interference in mosquitoes. In addition, these results provide a mechanistic understanding of the mode of action of *Wolbachia*-induced pathogen blocking in Anophelines, which is critical to evaluate the long-term efficacy of control strategies for malaria and *Anopheles*-transmitted arboviruses.

**HIGHLIGHTS:**

- *Wolbachia* inhibits O’nyong nyong virus (ONNV) in *Anopheles* mosquitoes.
- Enhanced Toll signaling is responsible for *Wolbachia*-induced interference of ONNV.
- Cholesterol suppresses Toll signaling to modulate *Wolbachia*-induced ONNV interference.

## INTRODUCTION

*Wolbachia*-based strategies to control arthropod-borne diseases are gaining considerable attention. After transinfection of *Wolbachia* into novel vectors (reviewed in Hughes and Rasgon, 2014), the bacterium often impairs the hosts ability to become infected with and transmit additional pathogens. While release of *Wolbachia*-infected mosquitoes to control arbovirus transmission is well underway (Hoffmann et al., 2011; Walker et al. 2011, Utarini et al. 2021, Collin et al. 2022, Indriani et al. 2023), the molecular mechanisms underpinning *Wolbachia*-meditated pathogen interference remain unclear. Two main theories have been postulated regarding the mechanism of *Wolbachia*-mediated pathogen interference; immune priming of the host by *Wolbachia* infection that subsequently impedes other pathogens, or competition between *Wolbachia* and pathogens for metabolic resources such as cholesterol (Frentiu 2017, Geoghegan et al., 2017). While evidence exists to support both hypotheses (Bian et al. 2010, Pan et al. 2012, Bourtzis et al., 2014; Rainey et al., 2014, Frentiu 2017, Geoghegan et al., 2017, Jiménez et al. 2021), unambiguous general support for either hypothesis is yet to be provided. In addition, interference mechanisms are likely to differ between insect hosts, *Wolbachia* strains, pathogens, and the type of association (natural or artificial). Given that *Wolbachia*-infected mosquitoes are currently being released into field populations (Hoffmann et al., 2011; Walker et al. 2011, Collin et al. 2022, Utarini et al. 2021, Indriani et al. 2023), there is a great urgency to understand the molecular mechanisms underpinning these pathogen interference phenotypes.

In support of the hypothesis that immune priming by *Wolbachia* subsequently impedes other pathogens, mosquitoes artificially infected with *Wolbachia* often have enhanced innate immunity (Bian et al., 2010; Pan et al. 2012, Blagrove et al., 2012; Kambris et al., 2009; Moreira et al., 2009, Hughes et al. 2011a, 2011b). For example, in *Aedes aegypti* infected with the *w*AlbB *Wolbachia* strain, elevated levels of reactive oxygen species (ROS) lead to upregulation of the Toll pathway, decreasing mosquito susceptibility to dengue virus (Pan et al., 2012). Similarly, *Anopheles stephensi* stably infected with *w*AlbB have higher ROS levels compared to their uninfected counterparts (Bian et al., 2013), suggesting that a similar mechanism might be occurring in Anopheline mosquitoes. Additionally, ROS can directly inhibit *Plasmodium* development in *Anopheles* (Luckhart et al., 2013; Molina-Cruz et al., 2008).

While enhanced host immunity contributes to pathogen interference in mosquitoes, this is likely not the sole mechanism by which *Wolbachia* inhibits pathogens. In native *Wolbachia* associations in *Drosophila* where *Wolbachia* does not alter basal immunity (Bourtzis et al., 2000; Xi et al., 2008), a protective effect against viral pathogens is still observed (Bourtzis et al., 2000; Hedges et al., 2008; Teixeira et al., 2008; Xi et al., 2008). For example, *Wolbachia* infection in *Drosophila melanogaster* mutants deficient in Toll and IMD signaling still resulted in impaired viral infections (Rances et al., 2013; Rances et al., 2012). These studies suggest that immune induction is not the sole mechanism by which *Wolbachia*-mediated pathogen protection occurs in Diptera. There is mounting evidence that resource competition between *Wolbachia* and other pathogens contributes to both pathogen interference (artificial associations that block pathogen transmission) and pathogen protection (natural associations that protect the host). In particular, competition for cholesterol, a common nutritional requirement for both *Wolbachia* and viral and protozoan pathogens, may modulate pathogen development. In mosquitoes, *Wolbachia* behaves as a cholesterol heterotroph, depleting hosts cholesterol levels (Caragata et al., 2013). *Wolbachia*-infected *Drosophila* that received cholesterol supplementation had higher viral titers and increased virus-induced mortality (Caragata et al., 2013, 2014). These observations suggest that cholesterol can negatively impact the protective effect of *Wolbachia* in insects; however, the molecular mechanisms behind this effect remain elusive. Understanding the role of cholesterol in *Wolbachia*-meditated pathogen interference is particularly critical in mosquitoes as cholesterol is a natural component of human blood and is therefore ingested as part of a blood meal.

Using the O’nyong nyong virus (ONNV) – somatic *Wolbachia* infection – *Anopheles* mosquito system, we show that *Wolbachia* infection significantly reduces ONNV levels in artificial transient *in vivo* and stable *in vitro* associations. We investigated the molecular mechanisms underpinning *Wolbachia*-mediated pathogen interference of ONNV in *Anopheles* and found that (1) Toll-based immunity is central to protection and (2) manipulation of cholesterol levels significantly modulates the effect of *Wolbachia* on viral titers as had been previously observed in *Drosophila*. Strikingly, the increase in viral titers was an effect of cholesterol-induced modulation of Toll signaling rather than nutritional competition between *Wolbachia* and ONNV. These findings provide a functional link between the two main current hypotheses regarding the molecular mechanisms of *Wolbachia*-mediated pathogen interference in mosquitoes and highlight the need for further research into the role of ingested human cholesterol (and potentially other factors) on the vector competence of *Wolbachia*-infected mosquitoes.

## RESULTS

### *Wolbachia* inhibits ONNV levels in cell lines and mosquitoes

To determine if *Wolbachia*-mediated pathogen interference occurs during ONNV infection in mosquitoes, we first challenged *An. gambiae* Sua5B cells which were stably infected with either *Wolbachia* strain *w*Mel from *D. melanogaster* or *Wolbachia* strain *w*AlbB from *Aedes albopictus* (Rasgon et al. 2006) (Figure 1A). Both strains of *Wolbachia* significantly inhibited infectious ONNV titers in the cells (Figure 1B). This effect was density-dependent as evidenced by the increase in ONNV titers following antibiotic treatment of *w*AlbB-infected Sua5B cells (Figures 1C).

**Fig. 1.**
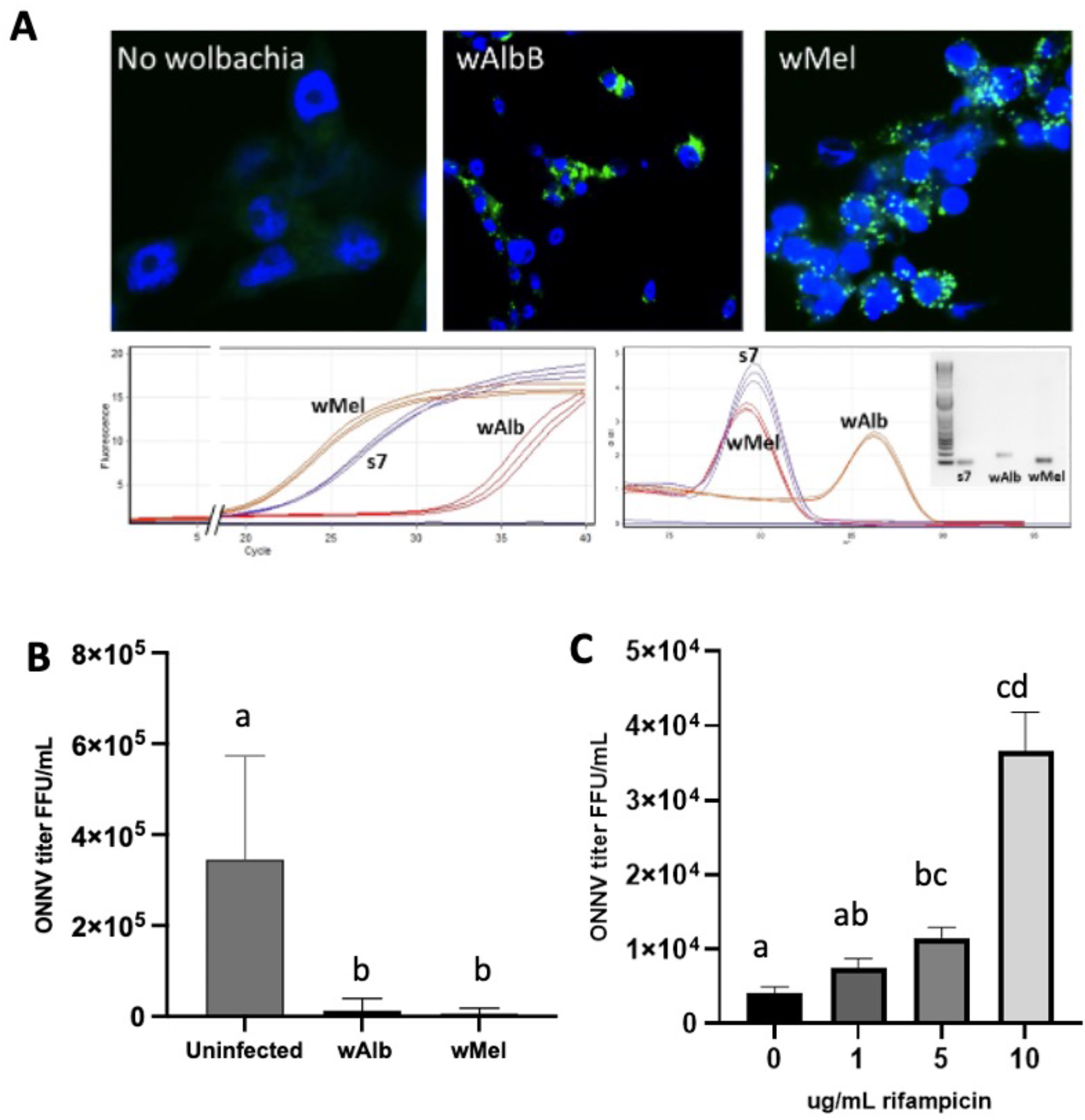
*Wolbachia* infection in *Anopheles gambiae* cells and inhibition of ONNV. A. Fluorescent *in situ* hybridization (FISH) of *Wolbachia* strains *w*Mel and *w*AlbB in infected Sua5B cells, gene-specific qPCR and melt curves, and resolution of PCR products on a gel, confirming cellular infection. B. Infectious ONNV titer in *Wolbachia* infected or uninfected Sua5B cell culture supernatants; both *Wolbachia* strains wAlbB and wMel significantly inhibit virus. C. ONNV is negatively correlated with *Wolbachia* density. As *Wolbachia* titers decrease due to antibiotic treatment, ONNV increases. Treatments with different letters are significantly different.

In order to confirm our results *in vivo*, we established transient somatic *Wolbachia* infections (*w*Mel and *w*AlbB) in female adult *An. gambiae* and *An. stephensi* mosquitoes by intrathoracic microinjection (Hughes et al., 2011b; Jin et al., 2009) and challenged mosquitoes 7 days later with an infectious blood meal containing ONNV. In both *An. gambiae and An. stephensi* we observed significant reductions in ONNV infectious viral titers in *Wolbachia*-infected individuals compared to uninfected controls (Figure 2). Given the similarities between the two different strains of *Wolbachia* in our experiments, further characterization studies were conducted with *w*AlbB.

**Fig. 2.**
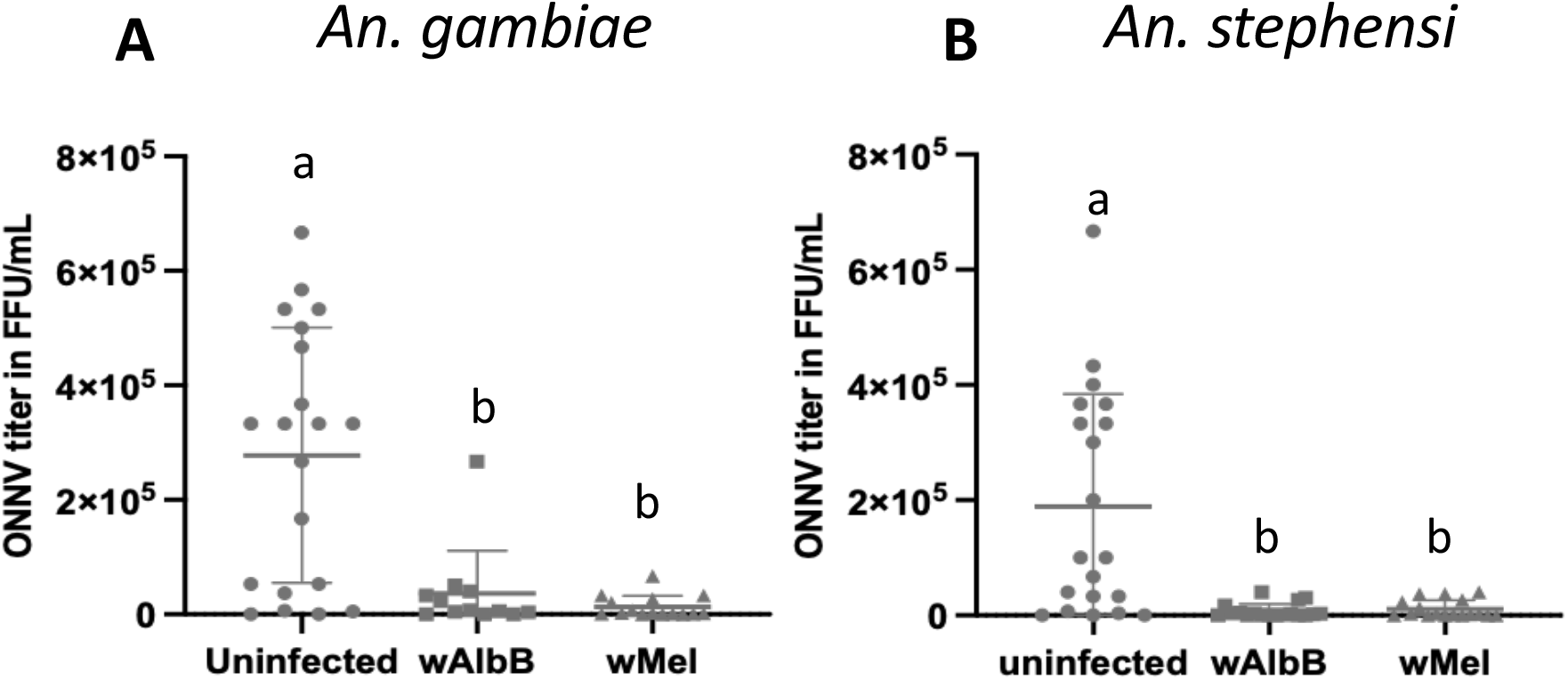
Transient somatic *Wolbachia* infections (wAlbB and wMel) inhibit ONNV in. A. *An. gambiae* and B. *An. stephensi* mosquitoes *in vivo*. Treatments with different letters are significantly different.

### *Wolbachia* upregulates *Anopheles* host genes antagonistic to ONNV

The Toll and IMD pathways are the main innate immune pathways in *Anopheles* mosquitoes (Christophides et al. 2002, Carissimo et al. 2014). Waldock and colleagues (2012) identified several *Anopheles* host genes which were antagonistic to ONNV, including genes in the Toll innate immunity signaling pathway such as *ML1* and *LYSC4*. As the Toll pathway has been shown to contribute to *Wolbachia* blocking of dengue virus in *Aedes aegypti* (Pan et al. 2012) we sought to determine if this immune pathway contributed to *Wolbachia*-mediated blocking of ONNV in *Anopheles*. We used RNA interference (RNAi) to knock down (KD) expression of key Toll and IMD pathway genes (including those identified by Waldock et al. to be involved in ONNV modulation) in *w*AlbB-infected *An. gambiae* cells, and quantified the effect on ONNV titers. We observed that KD of the Toll pathway genes *LYSC4*, *ML1*, *TOLL5A*, or the double KD of *ML1* and *TOLL5A* significantly increased ONNV titer in *Wolbachia*-infected cells (Figure 3A), indicating these genes influence the *Wolbachia* interference phenotype. No effect on ONNV titers was observed after KD of *IMD*, suggesting this pathway is not involved in the *Wolbachia* interference phenotype in *Anopheles* (Figure 3A). We further confirmed our results by reactivating the Toll pathway downstream of *ML1* and *TOLL5A* by KD of the negative regulator *cactus* in conjunction with KD of TOLL or ML1. KD of *cactus* restored the protective effect of *Wolbachia* against ONNV (Figure 3A).

**Fig. 3.**
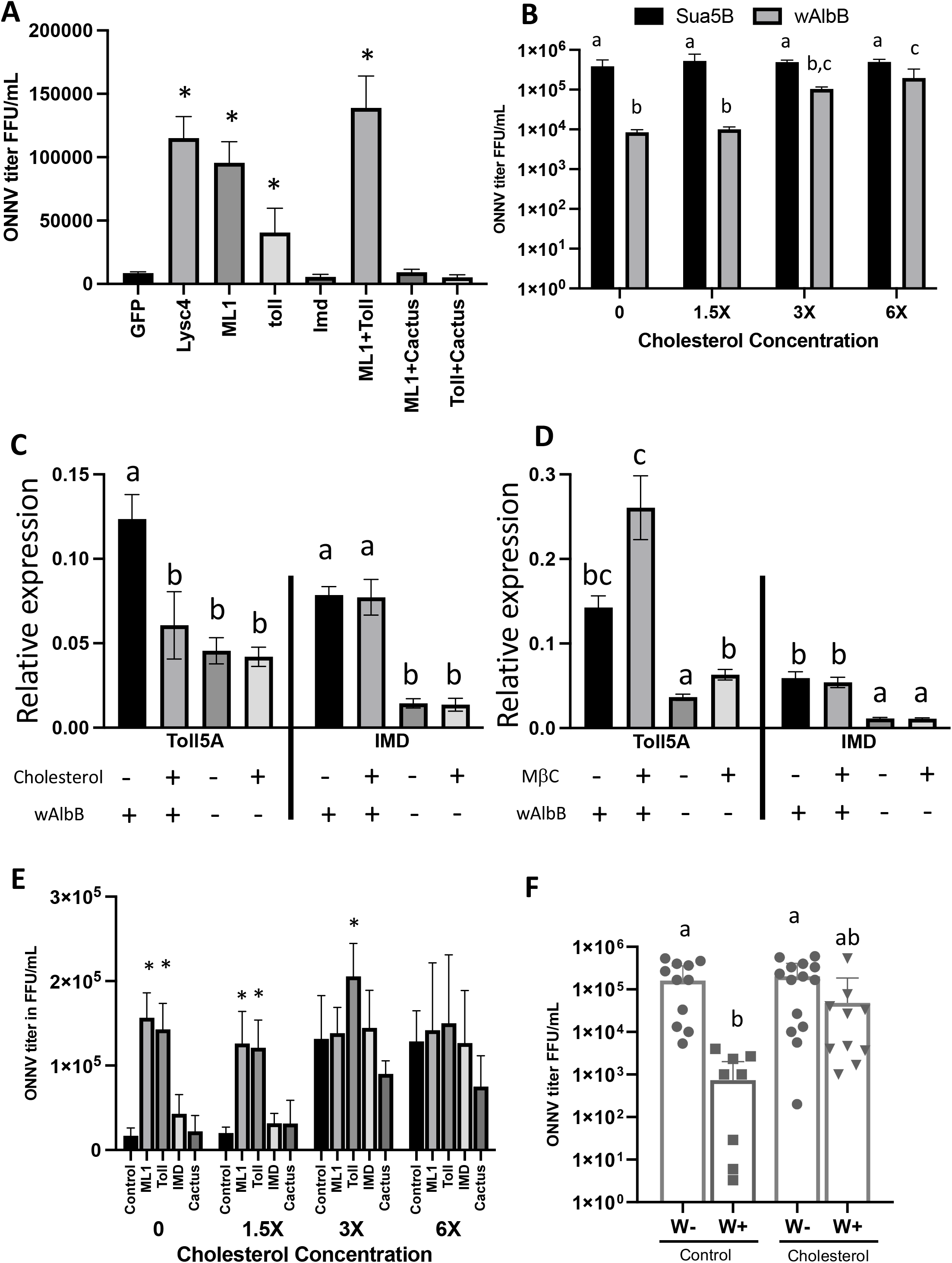
Interactions between *Wolbachia*, cholesterol, mosquito innate immunity, and ONNV. A. Effect of immune gene knock-down in wAlbB-infected cells on ONNV titers. Depletion of *LYSC4*, *ML1* or *TOLL5A* (or *ML1* + *TOLL5A* double KD) ablates *Wolbachia*-induced ONNV inhibition, while reactivation of the Toll pathway through *cactus* knock-down completely restores the inhibition phenotype. B. Cholesterol inhibits *Wolbachia*-induced ONNV inhibition in a dose-dependent manner, but has no effect on ONNV levels in *Wolbachia*-uninfected cells. C. Cholesterol supplementation suppresses *TOLL5A* expression in wAlbB-infected but not *Wolbachia*-uninfected cells. While *Wolbachia* induces *IMD*, there is no effect of cholesterol on *IMD* expression in *Wolbachia* infected or uninfected cells. D. Cholesterol sequestration induces *TOLL5A* expression in wAlbB-infected and uninfected cells, and the effect is synergistic with *Wolbachia*, but has no effect on IMD expression. E. Cholesterol supplementation eliminates *cactus*-based reactivation of Toll signaling and *Wolbachia*-mediated suppression of ONNV in a dose-dependent manner. F. Cholesterol supplementation partially ablates *Wolbachia*-induced inhibition of ONNV *in vivo* in *An. gambiae*. Treatments with different letters are significantly different.

### Cholesterol inhibits *Wolbachia*-induced activation of Toll signaling in ***Anopheles* cells**

To determine the effects of cholesterol supplementation on *Wolbachia*-mediated ONNV interference, we supplemented *w*AlbB-infected and uninfected cells with cholesterol and then challenged them with ONNV. Cholesterol supplementation did not significantly alter ONNV titer levels in *Wolbachia*-uninfected Sua5B cells. However, the protective effect of *Wolbachia*-infection against ONNV infection was lost in a dose-dependent manner following cholesterol supplementation (Figure 3B).

Because we had evidence for both TOLL-based innate immunity and cholesterol in mediating *Wolbachia*-induced pathogen blocking, we examined if there was a link between the two pathways. *Wolbachia* infection upregulated expression of *TOLL5A* (Figure 3C). We found that cholesterol supplementation inhibited *TOLL5A* expression in *Wolbachia*-infected cells, but had no significant effect in *Wolbachia*-uninfected cells (Figure 3C). Conversely, treatment with the cholesterol sequestering chemical methyl-beta-cyclodextrin (MßC) increased *TOLL5A* expression in both *Wolbachia*-infected cells and uninfected cells, but the effect was amplified by the presence of *Wolbachia* (Figure 3D). Neither cholesterol supplementation nor sequestration affected expression of *IMD* in either *Wolbachia*-infected or uninfected cells (Figures 3C, 3D).

To confirm the interplay between cholesterol and Toll-based immunity in *Anopheles*, we undertook RNAi KDs in combination with cholesterol supplementation in *w*AlbB-infected Sua5B cells. In the GFP control KD, cholesterol ablated the protective effect of *Wolbachia* in a dose-dependent manner. KD of *ML1* or *TOLL5A* uniformly increased ONNV titers in the no-cholesterol control. KD of *cactus* (which activates Toll signaling downstream of *TOLL5A* and *ML1*) restored the protective effect of *Wolbachia*, but as cholesterol levels increased this effect was lost in a dose-dependent manner. KD of *IMD* in the presence of cholesterol had no effect on ONNV levels (Fig 3E).

Finally, we confirmed this effect *in vivo* by feeding *An. gambiae* ONNV-infected blood with or without cholesterol. Supplementation of cholesterol in an ONNV infectious blood meal significantly reduced the protective phenotype of *Wolbachia* in *Anopheles* mosquitoes but had no effect in *Wolbachia*-uninfected mosquitoes (Figure 3F).

## DISCUSSION

### *Wolbachia*-based suppression of ONNV replication through Toll-mediated immunity is inhibited by cholesterol supplementation

We have shown that *Wolbachia* inhibits development of ONNV in *Anopheles* cells and mosquitoes, and demonstrated a link between the two main hypotheses explaining *Wolbachia*-induced inhibition of pathogens (immune priming vs. metabolic competition). As observed in other mosquito systems (Pan et al., 2011), *Wolbachia* infection induced the Toll pathway which provided protection against invading viruses. Similar to *Wolbachia*-mediated protection in *Drosophila*, cholesterol supplementation in *Anopheles* dramatically ablates *Wolbachia*-mediated viral interference. However, this effect seems to be due to repression of TOLL signaling rather than metabolic competition for cholesterol between *Wolbachia* and virus.

*Anopheles ML1* has been shown to interact with cytoplasmic actin to mediate phagocytosis and killing of pathogens (Sandiford et al., 2015). In mammals, the *ML1* homologue *MD2* is the co-receptor for *TLR-4*, where *TLR-4* and *MD2* form a heteroduplex for binding of LPS and initiation of the pathway (Akashi et al., 2003; da Silva Correia et al., 2001; da Silva Correia and Ulevitch, 2002; Muroi et al., 2002). In contrast, Toll signaling in *Drosophila* is initiated by a complex of TOLL and *spaetzle* (Alpar et al., 2018; Chowdhury et al., 2019). Our data suggest that in addition to its previously described role in pathogen phagocytosis, *Anopheles ML1* is likely also involved in Toll signaling, suggesting that Toll signaling in *Anopheles* may be more similar to mammalian systems than to *Drosophila*.

In mammals, cholesterol is transported through the blood stream by two types of carrier lipoproteins: low-density lipoprotein (LDL) and high-density lipoprotein (HDL). Although both types of lipoproteins bind and transport cholesterol, they have very different structures, functions, and immunomodulatory effects (Michelsen et al., 2004; Xu et al., 2001). High levels of LDL have been associated with diseases such as atherosclerosis and rheumatoid arthritis and oxidized LDL can actually enhance TLR expression to induce a pro-inflammatory immune response (Howell et al., 2011; Li et al., 2020; Xu et al., 2001). In contrast, HDL has been shown to be anti-inflammatory and high levels of HDL in the blood are associated with protection from cardiovascular disease (Ben-Aicha et al., 2020; Catapano et al., 2014; Yu et al., 2010). Specifically, HDL has been shown to impair Toll signaling in human macrophages (van der Vorst et al., 2017). This is thought to occur, in part, due to the influence of cholesterol availability on the formation of lipid rafts, which in turn can modulate the function of a variety of immune receptors including TLRs (Fessler and Parks, 2011; Varshney et al., 2016).

The main constituent of HDL, apolipoprotein A-I, is highly conserved across vertebrate and invertebrate species, as is its ability to modulate cholesterol levels (Bashtovyy et al., 2011; Collet et al., 1997). Lipid transport in most insects occurs via lipophorin (Lp), which are a class of HDL (Chino et al. 1982, Shapiro 1988, Atella et al. 1992), that contains three apolipoproteins (ApoI, ApoII, ApoIII). Lps are involved in the transport of lipids (like cholesterol) to and from the fat body, the insect equivalent of the human liver (Marinotti et al., 2006). Both Lp and the precursor to ApoI/II have been documented to alter the immune response of mosquitoes to pathogens and parasites (Cheon et al. 2006). Specifically, in *Ae. aegypti* expression of Lp and the Lp receptor (LpRfb) were shown to be up-regulated in response to *Plasmodium gallinaceum* infection and KD of Lp significantly decreased oocyst development (Cheon et al., 2006). In *Anopheles,* the precursor of ApoI and ApoII appears to facilitate *Plasmodium* ookinete invasion and oocyst development (Mendes et al., 2008). Similarly, KD of ApoIII of *An. gambiae* also significantly increased *Plasmodium* development in the midgut (Gupta et al., 2010). Therefore it is possible that ingested cholesterol bound to Lp could be inhibiting Toll signaling pathways in *Wolbachia*-infected mosquitoes in a manner similar to what has been previously described for HDL in humans.

While induced immunity does not appear to be important for *Wolbachia*-induced pathogen protection in *Drosophila* (Rancès et al., 2012, 2013), our data suggest immunity is a major driving force behind virus interference in *Anopheles*. Given that Toll-based immunity also influences *Wolbachia*-mediated pathogen interference in *Aedes aegypti* (Pan et al., 2012), this may suggest the mechanism behind *Wolbachia* pathogen interference differs according to the insect association. Differences could be attributed to pathogen interference (interference of pathogens transmitted by an insect vector) compared to pathogen protection (protection of the insect host from deleterious pathogens) (Hughes and Rasgon, 2012), variation in the mechanism behind *Wolbachia* strains (wAlbB compared to wMel-like strains), or host variation (mosquitoes compared to flies). Variation in the protective effect of different strains of *Wolbachia* has also been demonstrated (Chrostek et al., 2013).

From an applied perspective*, Wolbachia* is under investigation as a means to control malaria in *Anopheles* mosquitoes (Hughes et al., 2011b, Bian et al., 2013, Kambris et al. 2010, Jeffries et al. 2018, Walker et al. 2021). ONNV provides a tractable *in vitro* system to investigate *Wolbachia*-mediated pathogen protection in *Anopheles* mosquitoes, which may shed light on the mechanism involved in the inhibition of *Plasmodium.* For example, silencing experiments indicate *ML1* is antagonistic to *P. falciparum* yet is an agonist of *P. berghei* (Dong et al., 2006). Up-regulation of this gene by *w*AlbB may explain the inhibitory effect on *P. falciparum* (Bian et al., 2013; Hughes et al., 2011b) and enhancement of *P. berghei* (Hughes et al., 2012) in transiently *Wolbachia*-infected *Anopheles.* The findings that *Wolbachia* can be vertically transmitted in *Anopheles* after manipulation of the native microbiota, the successful transinfection of *An. stephensi*, and the identification of native *Wolbachia* strains in natural *Anopheles* populations all further enhance the prospects of applied *Wolbachia* strategies in *Anopheles* (Hughes et al., 2011b, Hughes and Rasgon 2014, Bian et al., 2013, Kambris et al. 2010, Jeffries et al. 2018, Walker et al. 2021, Quek et al. 2022). While issues surrounding *Wolbachia* pathogen enhancement in mosquitoes have been raised (Hughes et al., 2014; Hughes et al., 2012; Murdock et al., 2014; Zele et al., 2014), our results suggests this is not a concern for ONNV, and possibly for other viruses capable of being transmitted by *Anopheles*, such as Mayaro virus (Brustolin et al. 2018, Terradas et al. 2022).

## EXPERIMENTAL PROCEDURES

### Mosquito and *Wolbachia* strains

*An. gambiae* mosquitoes (Keele strain) and *An. stephensi* (Liston strain) were reared at 27°C and relative humidity of 85% with a 12-hour photoperiod and offered 10% sucrose *ad libitum*. The *w*AlbB and the *w*Mel strains of *Wolbachia* were cultured in Sua5B cells as previously described (Rasgon et al., 2006). Fluorescence *in situ* hybridization (FISH) visualization of endosymbionts and qPCR/PCR detection in cell lines was carried out as previously described (Rasgon et al. 2006, Hughes et al. 2011a, 2011b). To infect *Anopheles* mosquitoes with *Wolbachia* the bacterium was purified from cells lines (Rasgon et al., 2006) and 100nl *Wolbachia* suspension (10^8^ bacteria/mL) or medium (control) was intrathoracically microinjected into two day old anesthetized female *Anopheles* mosquitoes (Hughes et al., 2011b; Jin et al., 2009).

### ONNV production, infection of cells and quantification

ONNV was generated from the full-length ONNV infectious clone p5’dsONNic. RNA was *in vitro* transcribed from linearized *Not*I digested plasmid and purified using RNeasy kits (Qiagen). Two micrograms RNA was transfected into Vero cells using Lipofectamine LTX (Invitrogen) and grown as previously described (Pujhari et al. 2022). Infected cells were cultured at 32°C with 5% CO_2_ for 72 hours, then supernatant of the culture was harvested, stored at –80°C and used when required for viral infections of cells or mosquitoes. Virus stock titers were determined by FFU assays using Vero cells (Brault et al., 2004) as previously described (Pujhari et al. 2022, Terradas et al. 2022).

Sua5B cells with and without *Wolbachia* (Rasgon et al., 2006) were infected with ONNV to assess for *Wolbachia*-mediated pathogen interference. Cell lines were cultured in 24-well plates at 25°C until confluent, then cell culture media (Schneider’s with 10% FBS) was removed and replaced with 500 µl fresh media containing 1×10^7^ FFU of virus and incubated at room temperature with constant shaking for 2 hours. The media was then removed and replaced with fresh Schneider’s media with 10% FBS. Cells were cultured at 25°C for 5 days before ONNV density in the supernatant was assessed via FFU assay as previously described.

### *Wolbachia* density-dependence antibiotic assay

*w*AlbB-infected Sua5B cells were treated with rifampicin for 4 h at 10 µg/ml, 5 µg/ml or 1 µg/ml, respectively (Lu et al., 2012). After treatment, cells were left to recover for 24 hours then infected with ONNV, and ONNV in the supernatant quantified by FFU assay 5 days later as described above.

### ONNV and *Wolbachia* infection of mosquitoes

5 day-old *An. gambiae* and *An. stephensi* were infected with wAlbB or wMel by intrathoracic injection as previously described (Jin et al. 2009, Hughes et al. 2011b) and *Wolbachia* allowed to establish for 7 days. Age-matched *Wolbachia*-infected or uninfected (media injected) mosquitoes were orally infected with ONNV by an infectious blood meal using a membrane feeder. Prior to feeding, mosquitoes were starved overnight. 10^7^ FFU ONNV was presented to mosquitoes in blood warmed to 37°C for feeding. After blood feeding, unfed mosquitoes were removed. Six days post feeding, mosquitoes were harvested, homogenized individually and the lysate directly tested for infectious virus titer by FFU assay as described above.

### Immune gene expression and RNAi in cell lines

Candidate mosquito genes that affect ONNV (*ML1* and *Lysc4*) were selected from Waldock (2012) as well as canonical genes in the Toll and IMD signaling pathways (*TOLL5A*, *IMD*), and the negative regulator of Toll (*cactus*). Gene expression was normalized to the S7 gene. To determine if these genes were involved in the *Wolbachia*-mediated pathogen interference of ONNV, genes were knocked down by RNAi in *w*AlbB infected Sua5B cells. Gene-specific double stranded RNAs (dsRNAs) were synthesized using T7 MEGAscript kit (Ambion) as described (Waldock et al., 2012), and transfected into cell lines. ONNV infection of cell lines was performed 3 days post transfection and expression levels of the target genes cells checked by qPCR to confirm knock down (Supplementary Figure 1). Three days post ONNV infection, virus density was quantified in the culture supernatants by FFU assay as described above. All primer sequences are given in Supplementary Table 1.

### Effect of Cholesterol on ONNV and host gene expression in the context of ***Wolbachia* infection**

To examine the effect of cholesterol on *Wolbachi*a-mediated pathogen interference, cholesterol (250X Cholesterol lipid concentrate, #12531018, Thermo Fisher) was supplemented to *w*AlbB-infected and uninfected Sua5B cells (0, 1.5X, 3.0X, or 6.0X) and infected with ONNV as described above. Viral titers in the supernatants were quantified 3 days post-infection by FFU assay as described.

To examine the effect of cholesterol on candidate gene expression in *Wolbachia* infected and uninfected backgrounds, wAlbB-infected and uninfected cells were washed with FBS-free medium and incubated in FBS-free medium for 3h in 48-well plate. Cells were then incubated with 3X cholesterol or 5mM methyl beta cyclodextrin (MßC) (#377110050, Thermo Fisher) (doses chosen for biological effectiveness but minimal cellular toxicity [Supplementary Figures 2 and 3]) for 4 hours, washed, then incubated with Schneider’s media with 10% FBS at 25°C for 4 h prior to gene expression qPCR.

To assess the effect of cholesterol treatment on *Wolbachia* blocking of ONNV and the role of candidate gene expression, we supplemented wAlbB-infected cells with cholesterol (0, 1.5X, 3.0X, 6.0X) while simultaneously knocking down *MDL1*, *TOLL5A, IMD, or cactus* or (compared to *GFP* control) using dsRNA as described above. Cells were infected with ONNV 3 days post-cholesterol treatment and dsRNA treatment and virus titer in the culture supernatants quantified by FFU assay 3 days post-infection.

### Statistical analysis

Data were analyzed by Analysis of variance (ANOVA) with Tukey’s correction for multiple comparisons.

## AUTHOR CONTRIBUTIONS

Conceived the study: GLH, JLR. Performed the experiments: SP, GLH, YS. provided reagents: JLR. Analyzed the data: GLH, NP, JLR. Wrote the paper: SP, GLH, NP, JLR.

## ACKNOWLEDGEMENTS

We thank Dr. Brian Foy for providing the ONNV infectious clone, Dr. Christopher Cirimotich, Ping Xue, and Rhiannon Schneider for technical support, Dr. Long Cui for experimental assistance, and the Penn State Huck Institutes Microscopy and Cytometry Facility for assistance with microscopy. This research was supported by NIH grants R01AI116636, USDA Hatch funds (Project PEN04769), a grant with the Pennsylvania Department of Health using Tobacco Settlement Funds, and funds from the Dorothy Foehr Huck and J. Lloyd Huck endowment to JLR. GLH was supported by grants from the BBSRC (BB/V011278/1, BB/T001240/1, BB/X018024/1, and BB/W018446/1), the UKRI (20197), a Royal Society Wolfson Fellowship (RSWF\R1\180013), the NIHR (NIHR2000907), and the Bill and Melinda Gates Foundation (INV-048598).

## Supplementary Figure Legends

**Supplementary Table 1.**
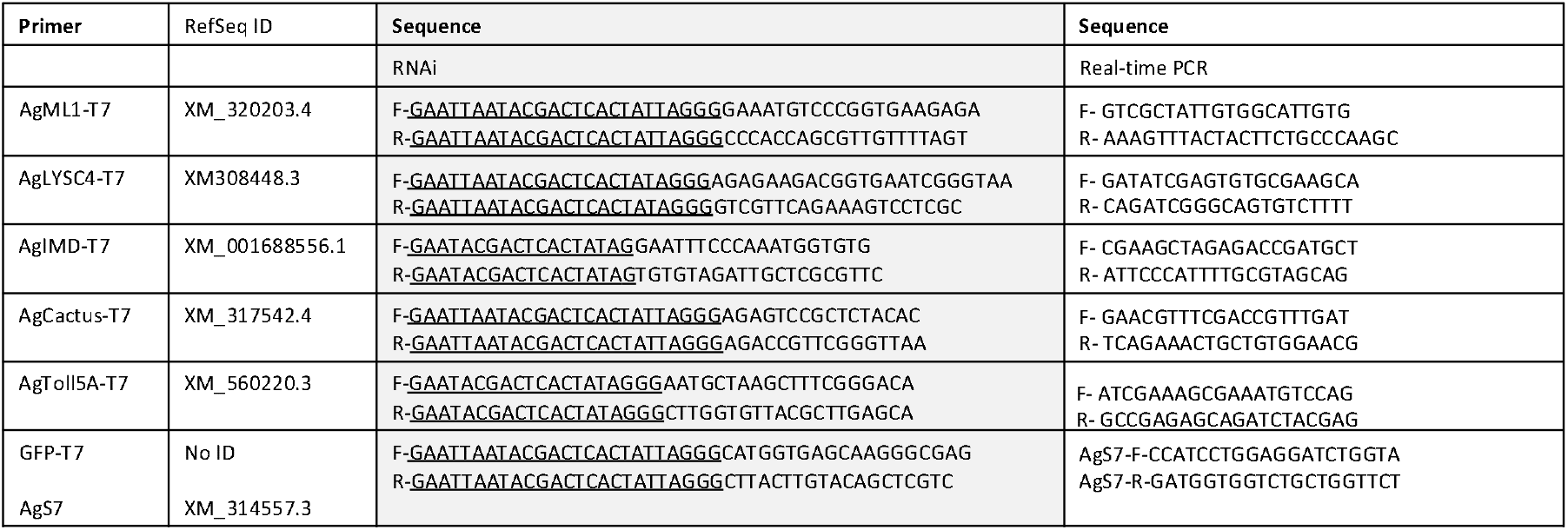
Primers used in this study.

**Figure S1.**
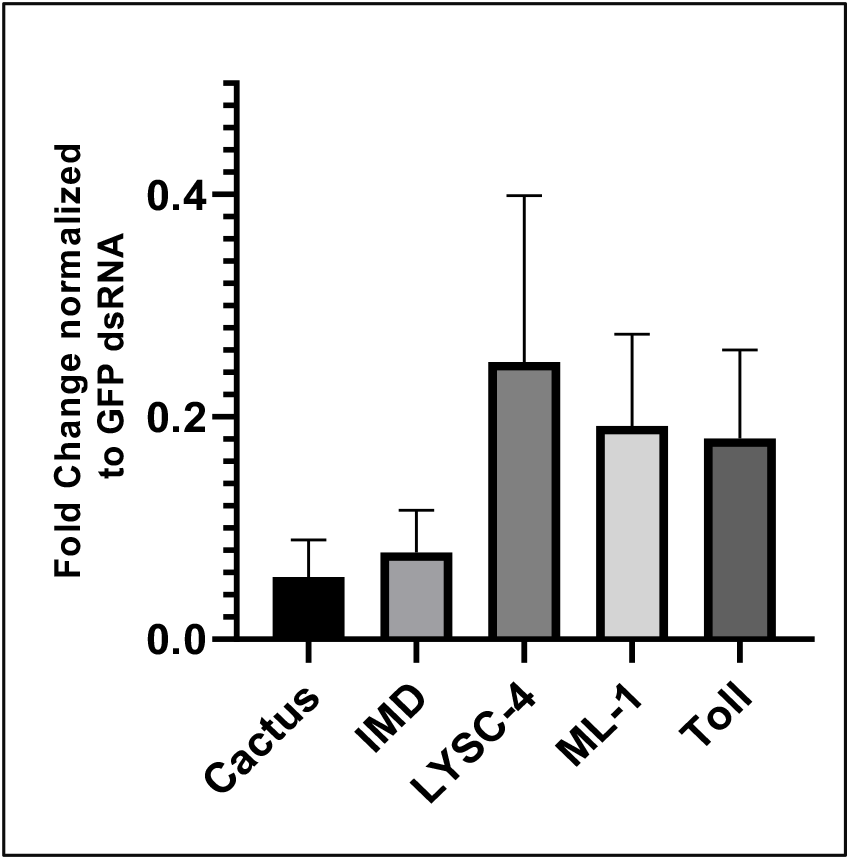
Confirmation of RNAi gene knockdown efficiency.

**Figure S2.**
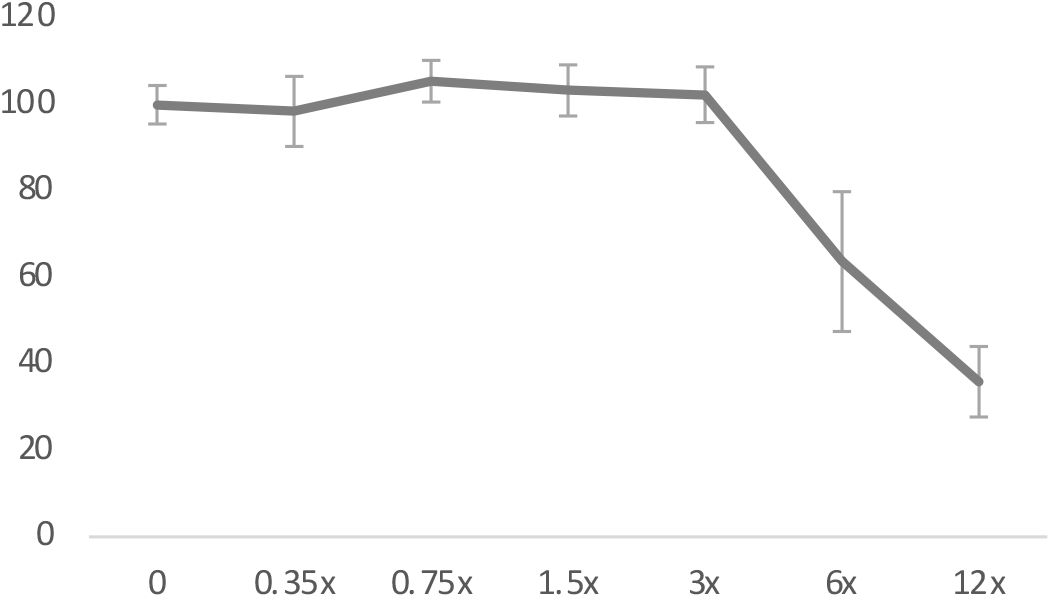
Dose-dependent cholesterol toxicity in Sua5B cells.

**Figure S3.**
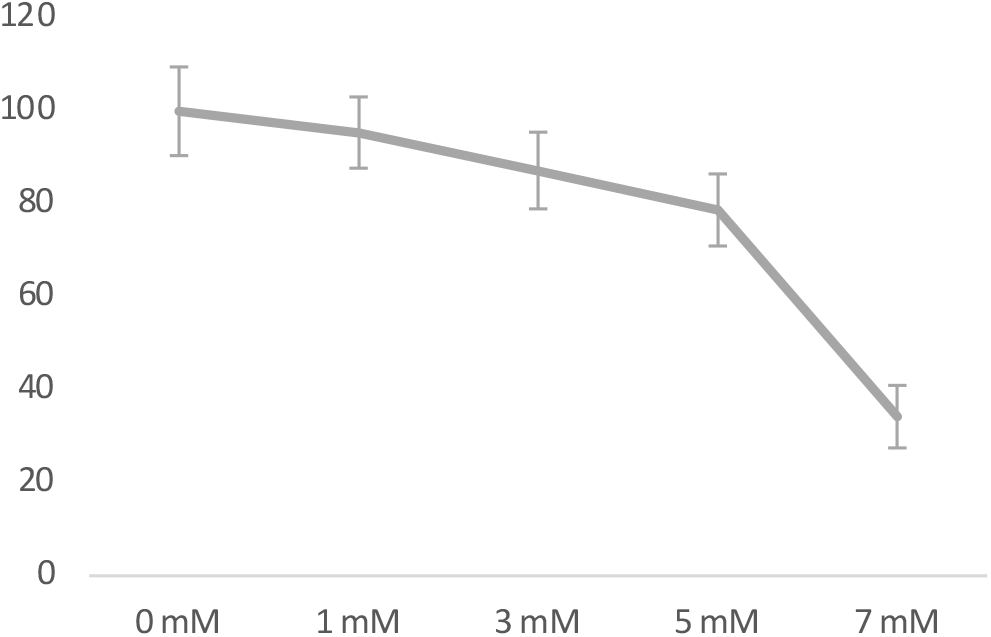
Dose-dependent methyl-beta-cyclodextrin toxicity in Sua5B cells.

## Notes

### Competing Interest Statement

The authors have declared no competing interest.

